# Unruly octopuses are the rule: *Octopus vulgaris* use multiple and individually variable strategies in an episodic-like memory task

**DOI:** 10.1101/2022.03.03.482865

**Authors:** Poncet Lisa, Desnous Coraline, Bellanger Cécile, Jozet-Alves Christelle

## Abstract

The evolution of complex cognition can be explained by different hypotheses, mutually non-exclusive: the social intelligence hypothesis, the ecological intelligence hypothesis and the predator-prey interaction hypothesis. Episodic-like memory can be used as a proxy to study complex cognition. This ability has mainly been studied through experimental tasks where subjects have to remember what they ate, where and when or in which context. Seemingly quite common in mammals and corvids, episodic-like memory abilities have been observed in only one invertebrate species: the common cuttlefish, a cephalopod mollusc. To explore if this ability is common to all cephalopods or if it has emerged to face specific constraints, we conducted an episodic-like memory task with seven *Octopus vulgaris*. Only one individual learnt the replenishing rates during the training and subsequently showed episodic-like memory abilities, whereas the other individuals favoured simpler foraging strategies, such as avoidance of familiarity and alternation, use of win-stay strategy and risk-sensitivity. A high variability in the use of these strategies was observed between and within individuals throughout the training. Since octopuses seem to live under lighter environmental pressure than cuttlefish, they may not need to rely on episodic-like memory abilities to optimize foraging as cuttlefish do. These results highlight the differences in the use of complex cognitive abilities between cuttlefish and octopuses, which might be linked with different environmental, predatory and social constraints.

**Summary statement:** When trained in an episodic-like memory task, common octopuses favour individually variable foraging strategies rather than keeping track of time to solve the task as cuttlefish do.

## Introduction

The evolution of complex cognition is currently explained by hypotheses emphasising either the importance of social or ecological challenges. The social intelligence hypothesis explains that advanced cognitive abilities evolved to answer the needs created by living in a group, such as maintaining social bonds, using cooperation and deception or learning from conspecifics (Byrne and Whiten, 1994; Dunbar, 1998). The ecological intelligence hypothesis supposes that complex cognition arose from the need to optimize foraging efficiency, such as manipulating food, finding diverse and multiple food types or track spatiotemporally dispersed food items (Byrne, 1997; Milton, 1981). Another ecological hypothesis is the predator-prey interaction hypothesis. It proposes that complex cognitive abilities might have emerged to cope with the challenges of finding and catching prey while avoiding predators (van der Bijl and Kolm, 2016). Evidence supporting social and ecological intelligence hypotheses mainly arise from studies undertaken in primates (Barrett et al., 2003; Rosati, 2017), mammalian carnivores (Holekamp et al., 2007), and birds (Emery et al., 2007). The predator-prey hypothesis finds its roots in various vertebrate species, such as fishes (Kotrschal et al., 2017; van der Bijl and Kolm, 2016), bovids (Köhler and Moyà-Solà, 2004) and is discussed with apes (Zuberbühler and Jenny, 2002). These evolutionary hypotheses are not necessarily mutually exclusive as: 1) variation of cognition might be the result of an intricate interaction between multiple selective pressures (Holekamp 2007); 2) specific cognitive skills might be an adaptation to cope with specific social and/or ecological challenges (Byrne and Bates, 2007; Rosati, 2017). Exploring the predictions of these hypotheses in invertebrates will be an important step to better understand main factors driving the evolution of cognition in distant phyla.

Different behaviours can be used as proxies for measuring complex cognitive abilities, and episodic-like memory is one of them. Episodic-like memory is the ability of an animal to remember the content (“what”), the spatiotemporal context (“where”, and “when” or “which”) and the source (contextual details such as the sensory modality of the content, the emotional valence, etc.) of a single event (Clayton et al., 2003). The ability to remember, in an integrated manner, the what, where and when (how long time ago) of an event has been shown in several taxa, including corvids (Clayton and Dickinson, 1998; Zinkivskay et al., 2009), rodents (sBabb and Crystal 2005) and apes (Ban et al., 2014; Martin-Ordas et al., 2010). Amongst invertebrates, common cuttlefish are, to our knowledge, the only species demonstrating episodic-like memory abilities (Jozet-Alves et al., 2013). In this experiment, cuttlefish ability to remember what they ate (shrimp or crab), where (position of the target) and how long ago (one or three hours) was tested. Identical targets at distinct locations (unique locations on each day) were associated with each prey type. Whereas the less preferred crab supply was replenished after any delay, the preferred shrimp supply replenished only after a long delay (three hours). Cuttlefish quickly learnt to go to the target delivering the preferred shrimp after a long but not after a short delay. Cuttlefish showed great capacities for the task, understanding the rules of the task in about 20 tri als (Jozet-Alves et al., 2013). A subsequent study confirmed the impressive memory abilities of cuttlefish, showing that their episodic-like memory does not fade, even in old age (Schnell et al., 2021). Another recent study showed that cuttlefish possess the ability to retrieve the sensory modality (seeing or smelling a prey) of a past event (Billard et al., 2020b), indicating that cuttlefish can bind the source of a memory in addition to remembering the content and the spatiotemporal context of their memory.

Modern cephalopods (i.e. octopods, cuttlefishes, and squids) are known for their complex cognitive abilities. Octopuses are generally considered as the most solitary (even if high-density populations have been reported; Godfrey-Smith and Lawrence, 2012; O’Brien et al., 2021; Scheel et al., 2017), squids as the most gregarious and cuttlefish in between (Boal, 2006). This suggests that complex cognition can arise even in the absence of strong social constraints. Their life is ruled by the need of eating without being eaten themselves. The lack of shell on a soft-bodied animal paired with the predatory pressure they live under may have forced cephalopods to develop efficient strategies relying on flexible behaviour and quickness to adapt, in other words complex cognitive abilities (Amodio et al., 2019), as stated in the predator-prey interaction hypothesis. Moreover, their constant need for food as indicated by their exponential growth (Hanlon and Messenger, 2018) may have promoted the development of efficient foraging strategies and memorization abilities to remember how, where and when to find food, enhancing their cognitive abilities as proposed in the ecological intelligence hypothesis.

We can wonder why cuttlefish possess episodic-like memory abilities. The first hypothesis is that this ability is shared with other large-brained cephalopod species as the result of their shared phylogeny. The second hypothesis is that episodic-like memory has emerged in cuttlefish to cope with specific ecological challenges: cuttlefish have to be constantly aware of predators while hunting, which requires time and energy and thus impact fitness. Their preys are often spatiotemporally dispersed in patches which often do not offer shelters, and when cuttlefish cannot minimize their risks by hunting from a hide, they use an array of cognitive skills to find prey at the right place and time, such as spatial memory (Jozet-alves et al. 2014), value-based decision making (Kuo and Chiao, 2020) or overcoming immediate gratification in order to obtain better preys (Schnell et al. 2021). Finally, the third hypothesis is that episodic-like memory in cuttlefish could be a mere by-product of the evolution of its complex cognition. It would have emerged from other abilities required by the cuttlefish to hunt and avoid predators, without any peculiar need for episodic-like memory itself.

Octopuses are also known for their impressive cognitive abilities. They live in a similar environment than cuttlefish, as they are shallow depth bottom dwellers (Hanlon and Messenger, 2018), but possess different means to handle their environmental constraints. Indeed, due to their lack of internal shell and their highly prehensible arms, octopuses possess a wider range of means of defence. While cuttlefish mainly use crypsis for defence, octopuses can also hide in crevices, arrange a shelter, cover themselves in rocks and shells in order to avoid attacks, or defend themselves aggressively against predators (Hanlon and Messenger, 2018). Moreover, thanks to their complex arms, octopuses’ food diet is broader than cuttlefish’s as octopuses can consume bivalves and gastropods, in addition to decapods, fishes and other cephalopods (Anderson et al., 2008; Mather et al., 2012).

Cuttlefish hunt moving preys living in patches such as crabs and shrimps which may come back to suitable patches quickly, replenishing the available preys promptly after being depleted. Remembering what was eaten where and when might thus be useful for cuttlefish. Octopus, on other hand, forage partly on sessile preys such as bivalves, which replenish on very long timescales, making the track of when places were visited unnecessary. Moreover, since octopuses possess multiple means of defence against predators, octopuses can wander more easily in the open compared to cuttlefish, finding preys as they go rather than relying on complex strategies such as the ones probably used by cuttlefish to minimize the time spent out of safety.

To optimize foraging efficiency, octopuses might rely on simpler foraging strategies based on rules of thumb which do not require a heavy cognitive load, such as the ones based on familiarity. Familiarity is a memory process which use a signal-detection function whereby elements exceeding a fixed criterion are recognized as having been perceived before (Yonelinas, 2001). The strategies which could be used include: a) spontaneous alternation, b) risk avoidance, and c) win-stay/win-shift strategies (Levine, 1959). a) Spontaneous alternation behaviour is the tendency to explore places that have been least recently explored (Ramey et al., 2009). In an experimental setting, the subject alternates the position of its choices on consecutive trials (Levine, 1959). Alternation is advantageous during exploration and foraging, especially when searching for preys like bivalves which replete slowly and represent a significant part of octopuses’ diet (Anderson et al., 2008). b) Risk-sensitivity is the foragers’ response to variance in food reward rate when choosing what to eat (Young et al., 1990). A food reward always available but of less quality will be considered less risky than a food reward of higher quality but randomly available. Some octopuses in the wild were observed favouring preys which were hard to find but presumably of higher nutritional or tasting quality, whereas other individuals did not strongly select their food and their choices followed prey availability (Mather et al., 2012). c) Win-stay-Lose-shift or Win-shift-Lose-stay strategies are used when subjects either repeat (stay) or avoid (shift) their last choice, depending on whether the choice was previously rewarded (win) or not rewarded (lose) (Kamil, 1983). Octopuses in the wild seem to use win-shift strategy when foraging, as they avoid returning to the same spot where they recently hunt, averting depleted locations (Mather 2014).

In order to assess whether octopuses keep track of time when different food sources vary in space and time or whether they favour simpler foraging strategies, we first evaluated their ability to learn replenishing rates of preferred versus less-preferred food items (procedure adapted from Jozet-Alves et al., 2013). Octopuses succeeding this task were subsequently tested to assess their episodic-like memory abilities (what-when-where experiment, adapted from Jozet-Alves et al., 2013). Our hypothesis was that given octopuses’ ecology, they might favour simpler foraging strategies based on familiarity, such as position preference, spontaneous alternation, risk sensitivity, and win-stay/win-shift strategies, rather than relying on time tracking strategies as cuttlefish do.

## Methods

### Ethical statement

Experiments were conducted in accordance with directive 2010/63/EU (European parliament) and with the French regulation applied to the protection and use of animals in research experiments. Procedures were approved (#22429 2019101417389263 v2) by the ethical committee of Normandy region (Comité d’Ethique de NOrmandie en Matière d’EXpérimentation Animale, CENOMEXA; agreement number 54).

### Subjects

The subjects used in the experiments were sub-adult common octopuses (*Octopus vulgaris* Cuvier). Octopuses were collected in the Mediterranean Sea by specialized fishermen (Carrodano, Poissons vivants, La Ciotat, France) in September 2020 (batch 1, n=3) and January 2021 (batch 2, n=4) (see Table 1 for names and sex). They were transported to the marine station of the University of Caen (Centre de Recherche en Environnement Côtier, Luc-sur-Mer, France). They were individually housed firstly in glass tanks of 50×50×50 cm and transferred in glass tanks of 100×50×50cm or 120×40×50cm as they grew. Octopuses were maintained in circulated semi-artificial seawater (salinity: 37 g/L, Instant Ocean Salt – Aquarium systems; temperature: 17±1°C; 7.8<pH<8.2; [NH_3_ + NH_4_ ^+^] < 0,25 mg/L; [NO_2_] < 0,2 mg/L; [NO_3_] < 50 mg/L), with artificial lighting following the natural light cycle. A sand bed, pebbles, shells and a shelter in the form of a terracotta pot or a PVC tube were provided in each tank. Octopuses were fed daily outside of the experimental trials with live crabs (*Hemigrapsus sanguineus* or *Carcinus maenas*), thawed or live shrimps (*Crangon crangon*), pieces of thawed fish (mackerel *Scomber scombrus*, pollock *Theragra chalcogramma*, herring *Clupea harengus* and whiting *Merlangius merlangu*s). Mussels (*Mytilus edulis*) were always available in the home tanks.

**Table 1:** Results of the food preference test for each octopus. Numbers within brackets correspond to the number of times a prey item was chosen during the test. Asterisks indicate a significant preference for one of the two prey types (binomial test, * p<0.05, ** p<0.01).

Figures
| Subject | Suricate | Abe | Pipoune | Coquille | Rosy | Tickle | Teddy |
| --- | --- | --- | --- | --- | --- | --- | --- |
| Batch | 1 | 1 | 1 | 2 | 2 | 2 | 2 |
| Sex | Male | Female | Female | Female | Female | Female | Female |
| Preferred prey | Crab (10) | Crab (10) | Crab (11) | Crab (10) | Crab (11) | Crab (10) | Crab (11) |
| Less preferred prey | Whiting (2) | Shelled mussel (2) | Mackerel (1) | Shrimp (2) | Shrimp (1) | Pollock (2) | Shrimp (1) |
| Binomial test, p-value | 0.039 * | 0.039 * | 0.006 ** | 0.039 * | 0.006 ** | 0.039 * | 0.06 ** |

### Procedure

Experiments were conducted in the home tank of each animal. Octopuses were pre-trained and tested for food preference, before starting the replenishing rate training (see Supplementary Materials for details).

#### Replenishing rate training

Octopuses were trained to learn that two different prey types (preferred *versus* less-preferred prey types; determined for each individual during the prey preference test) were available at specific locations and after specific delays (1h or 3h delay; Fig 1). Octopuses were tested five days a week: one trial per day, each trial consisting in two presentations separated by either a short (one hour) or a long (three hours) delay. During each presentation, octopuses were simultaneously presented with two closed opaque pots. Each pot contained a different prey item. The position and the content of the two pots were kept the same throughout the trials (“where” and “what” components were fixed for an individual for all the replenishing rate training).

**Figure 1:**
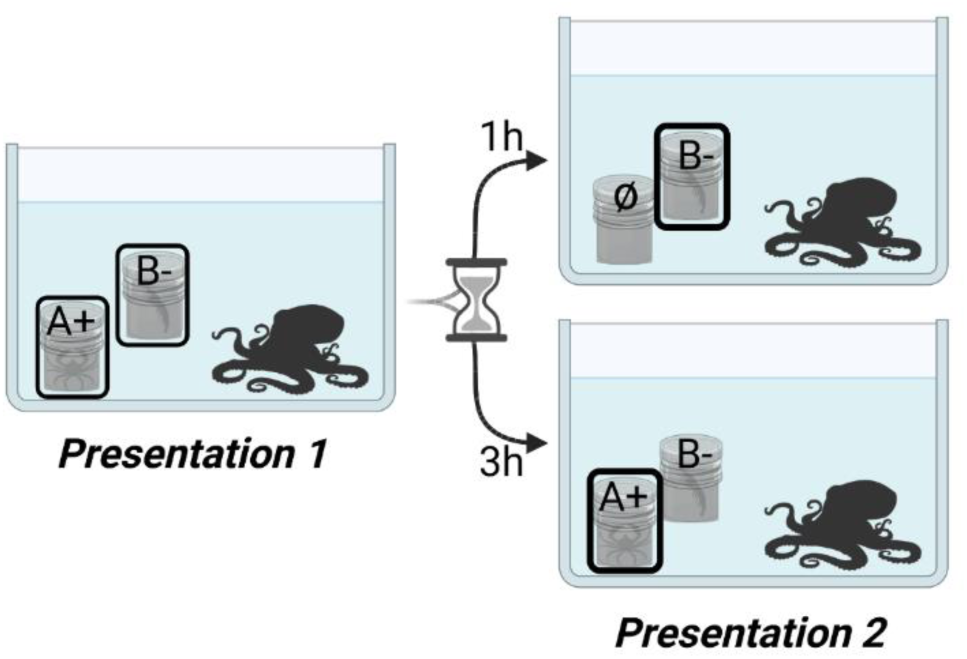
Organization of a training trial. During the presentation 1 of each trial, octopuses open both opaque pots to consume their preferred food (A^+^) and their less-preferred food (B^-^). After a one-hour delay, the pot previously containing A^+^ is empty, thus octopuses must go to the pot containing B^-^ to realize a successful choice. After a three-hours delay, both pots are replenished, and choosing the pot containing A^+^ was considered a successful choice. The position of the pots remained unchanged within trials. The position was altered between trials for the episodic-like memory task, but not for the replenishing rate task.

During the first presentation of a trial, the octopus could open and consume the content of each pot, and the pots were removed after 30 minutes. During the second presentation, pots were replenished according to the elapsed time since the first presentation. Following a short delay (1h), only the pot containing the less-preferred food item was replenished.

Following a long delay (3h), both pots were replenished. The octopus could only consume the content of one pot, the second pot being removed with a small net right after the choice. A choice was considered correct when an individual chose the pot containing the less-preferred food after a short delay, and the pot containing the preferred food after a long delay. An acquisition criterion was fixed at eight correct choices out of ten consecutive trials.

The maximum number of training trials was set to 40, corresponding to the double of trials cuttlefish needed to reach the acquisition criterion during previously published experiments (Jozet-Alves et al., 2013; cuttlefish learnt the replenishing rate in 21±4 trials). However, since the first batch of octopuses (n=3) did not reach the criterion in 40 trials, a second batch of octopuses (n=4) was subsequently constituted and the maximum number of training trials was set at 80.

#### Episodic-like memory task

Individuals which reached the acquisition criterion of the replenishing rate training were tested in the episodic-like memory task. This task was similar to the replenishing rate training task, excepting that the position of the pots changed each trial, while staying the same across the two presentations of a trial. During each trial, octopuses had to remember what prey was in each pot (what-where) and what time elapsed since the first presentation: spatiotemporal information was thus unique. We considered that octopuses showed episodic-like memory ability when they realized ten correct choices out of twelve consecutive trials (binomial test, p=0.039), with the maximum number of trials sets at 40 trials.

### Analysis

Data were analysed using R software (v. 3.5.1), using binomial tests for food preference tests and choices of octopuses. We first analysed choices of both batches of octopuses during the first 40 trials of replenishing rate training. We focused on the pots chosen during the second presentation to determine foraging strategies (replenishing rate use, familiarity, spontaneous alternation, risk sensitivity, Win-stay/Win-shift strategies) used by all individuals or by each individual separately. We compared choices during the first 20 and the last 20 trials of training, focusing only on the first 40 trials of replenishing rate training for both batches, and also explored the regularity of choices of individuals throughout the 40 trials of training. Two-tailed exact Fisher tests were used to compare the use of one strategy between the first and the last 20 trials of training. It should be noted that for alternation and Win-shift/Win-stay strategies, the choice on the first trial was excluded of the analyses, since there was no previous reference trial. Therefore, we analysed 39 trials and compared the first 20 trials with the last 19 trials of training for these strategies. To simplify the understanding of the following sections, we will use the expressions “40 trials” and “first and last 20 trials” as a way of speaking for all strategies.

## Results

### Food preference

All octopuses presented a significant preference for crabs (binomial test, p<0.039; Table 1). Less-preferred preys varied between individuals, with some octopuses tested with thawed fishes (whiting, mackerel or pollock), others with fresh shrimps or shelled mussels.

### Replenishing rate training and episodic-like memory task

In the first batch (maximum number of training trials sets at 40), none of the three octopuses reached the established learning criterion (*i*.*e*. eight correct choices out of ten consecutive trials). In the second batch (maximum number of training trials sets at 80), only one individual (Teddy) out of four reached the learning criterion in 43 trials (Fig. 2). Consequently, only Teddy was subsequently tested in the episodic-like memory task. The criterion for the task was set at 10 correct choices out of 12 consecutive trials (binomial test, p=0.039). Teddy reached the acquisition criterion in 21 trials.

**Figure 2:**
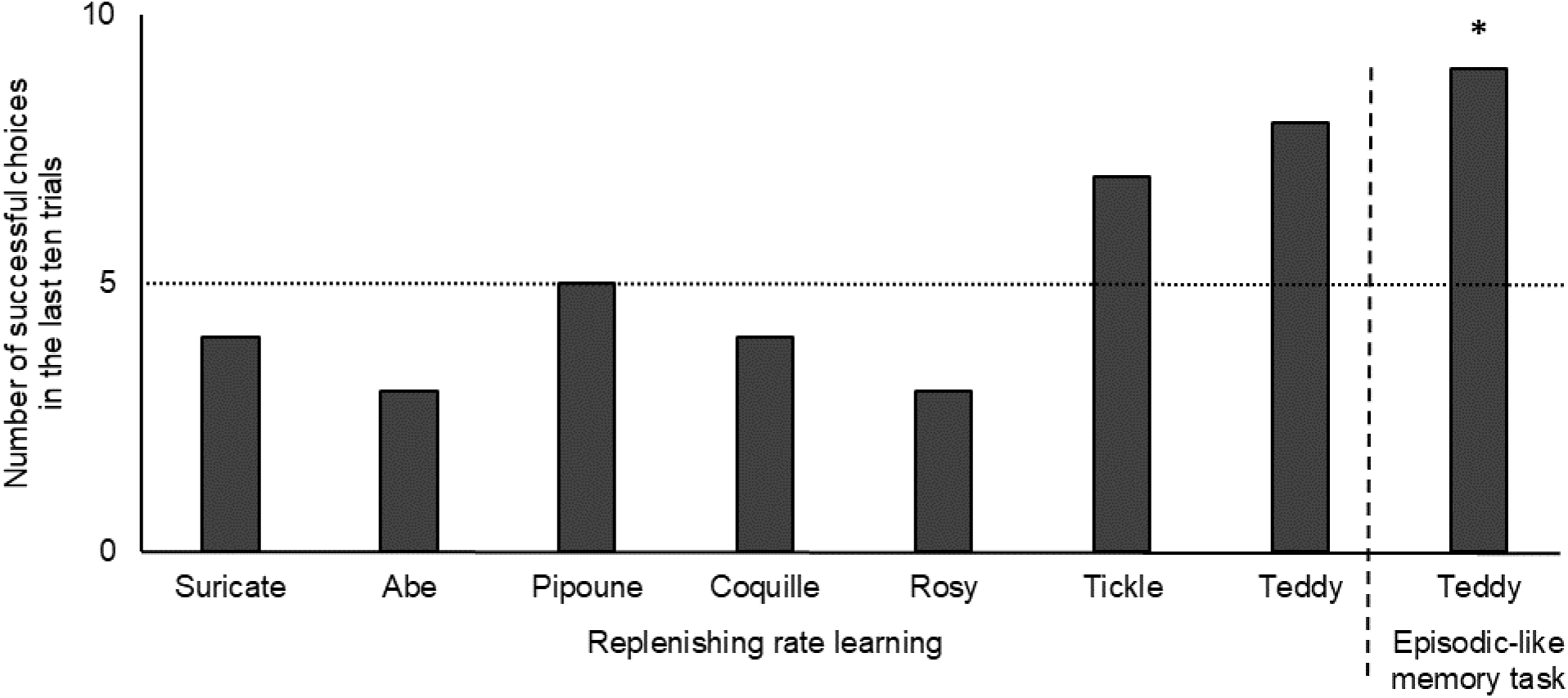
Number of successful choices in the last ten trials of the replenishing rate training and the episodic-like memory task. The asterisk represents a number of success significantly different from chance (i.e. dotted line; binomial test, * p<0.05).

### Strategies

Four strategies were analysed for the first 40 trials of training, for each octopus of both batches: familiarity, spontaneous alternation, risk-sensitivity and win-stay strategy (Fig. 3 for global/group analyses per strategy and Table 2 for individual analyses).

**Figure 3:**
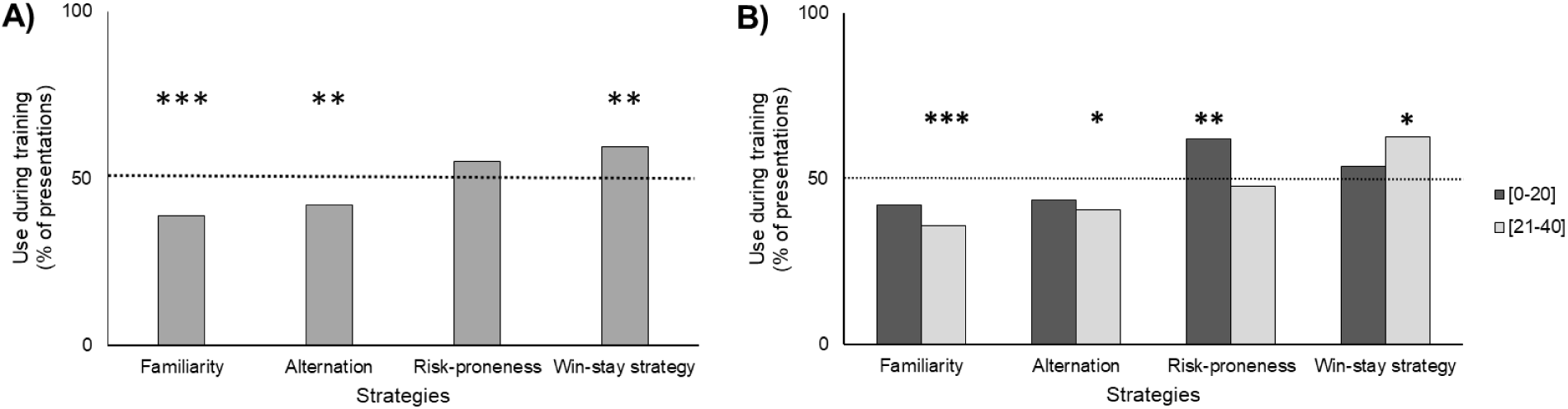
Percentage of octopuses (*n*=7) using each strategy during the replenishing rate training. A) Use of each strategy throughout the training. B) Use of each strategy throughout the training divided between the first and the last 20 trials. Asterisks represent significant difference from chance (*i*.*e*. dotted line; binomial test, * p<0.05, ** p<0.01, *** p<0.001).

**Table 2:** Summary of the strategies used by each individual. For familiarity, delays between brackets indicate a significant use of the strategy only after the delay specified. For risk-sensitivity, numbers between brackets indicate during which part of training the strategy was used ([2-21] for first 20 trials, [22-40] for the last 19 trials of training)

| Individual<br>Strategy | Suricate | Abe | Pipoune | Coquille | Rosy | Tickle | Teddy |
| --- | --- | --- | --- | --- | --- | --- | --- |
| Familiarity |  | Familiarity avoidance (3h) | Familiarity avoidance (3h) | Familiarity avoidance |  | Familiarity avoidance (1h) |  |
| Alternation | Constancy | Constancy | Constancy | Alternation | Constancy |  | Constancy |
| Risk-sensitivity | Averse [22-40] | Prone [2-21] [22-40] | Prone [2-21] Averse [22-40] | Prone [2-21] |  | Averse | Prone [2-21] |
| Win-stay | Win-stay |  |  |  |  |  | Win-stay |

#### Familiarity

Use of familiarity was observed when, during the second presentation of a trial, the subject chose the most familiar pot, which was in the place of the pot which was the last opened during the first presentation (also known as the second one opened during the first presentation; Fig. 4A).

**Figure 4:**
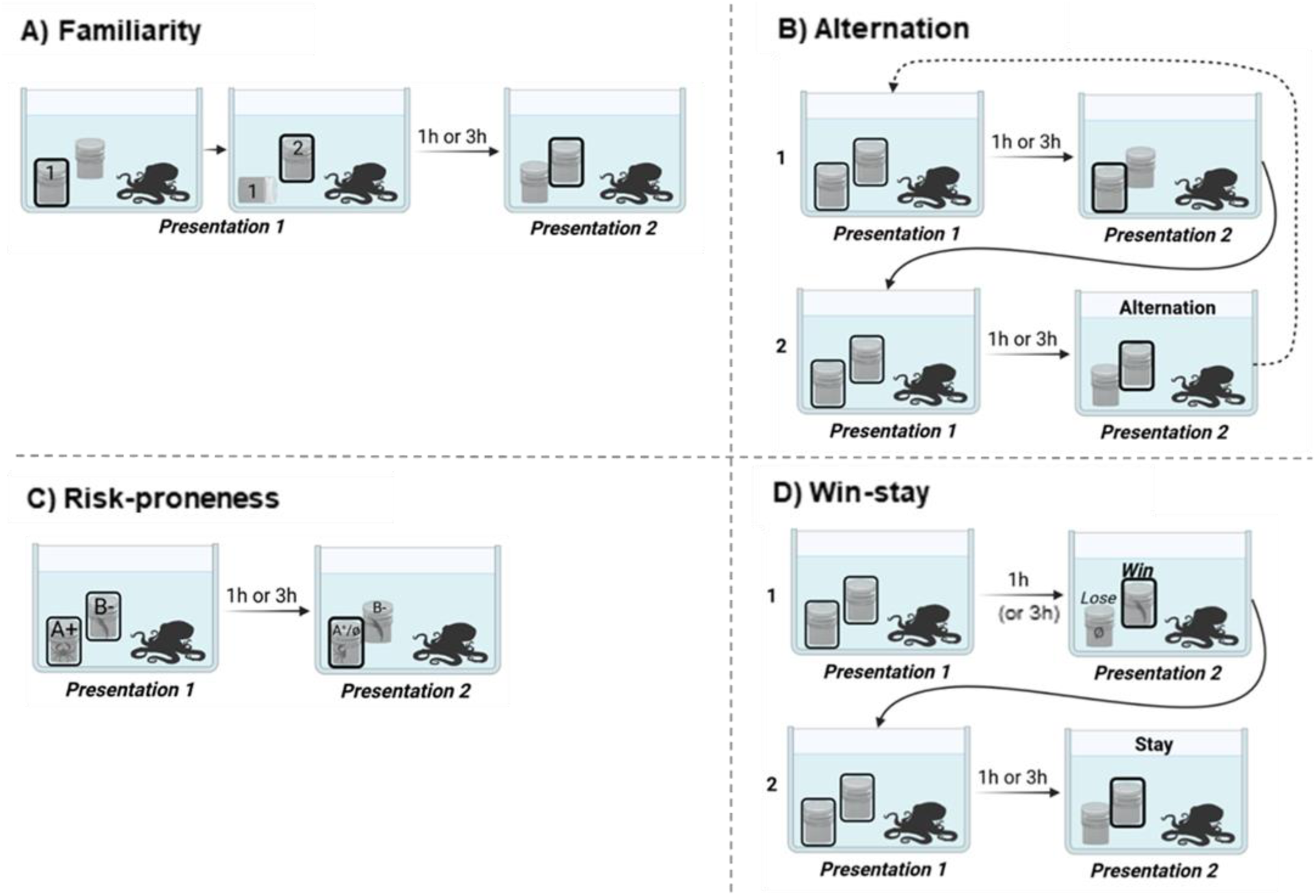
Representation of strategies used by octopuses during the replenishing rate training. A) Use of familiarity. During the presentation 1, octopuses choose a first pot, then a second one. On the presentation 2, they choose the pot lastly visited. B) Use of alternation. Octopuses choose one pot on the presentation 2 of a trial (1), and then choose the opposite pot on the presentation 2 on the following trial (2). C) Use of risk-proneness. During presentation 2, the pot containing the less-preferred food is less risky (B-) than the pot containing the preferred food (A^+^/Ø), since the less-preferred food is always available whereas the preferred food is available randomly if delays cannot be discriminated. D) Use of a Win-stay strategy. On the presentation 2, pots can either be a “win”, when replenished, or a “lose”, when empty. When octopuses open a pot with food inside on the second presentation during a trial (1; “win”), then on the second presentation on the following trial (2) they chose the same pot as the previous trial (“stay”), they use Win-stay strategy.

Considering all trials of all individuals, octopuses did not significantly use familiarity (109 familiarity choices out of 280 presentations, binomial test, p<0.001).

When distinguishing the first and the last 20 trials of training, octopuses significantly avoided the familiar pot during the last 20 trials of training (50/140, binomial test, p<0.001), but not during the first 20 trials of training (59/140, binomial test, p=0.076). However, the comparison between the first and the last 20 trials was not significant (two-tailed exact Fisher test, p= 0.499).

At the individual level, three octopuses (Tickle, Abe, Pipoune) significantly avoided the familiar pot all along the test (respectively 13/40 - 12/40 - 13/40, binomial test, p<0,039; Fig. 5A), and two individuals (Tickle, Coquille) avoided it nine times in a row during the last ten trials of training (binomial test, p=0.021). Among the three individuals significantly avoiding the familiar pot, it appeared that this behaviour was delay-dependent. Tickle significantly avoided the familiar pot after a short delay (4/20, binomial test, p=0.012), but not after a long delay (9/20, binomial test, p=0.012), whereas we observed the opposite behaviour in Abe and Pipoune with the familiar pot avoided after a long delay (respectively 3/20 - 3/20, binomial test, p=0.003), but not after a short delay (9/20 - 10/20, binomial test, p>0.824; Fig. 5B).

**Figure 5:**
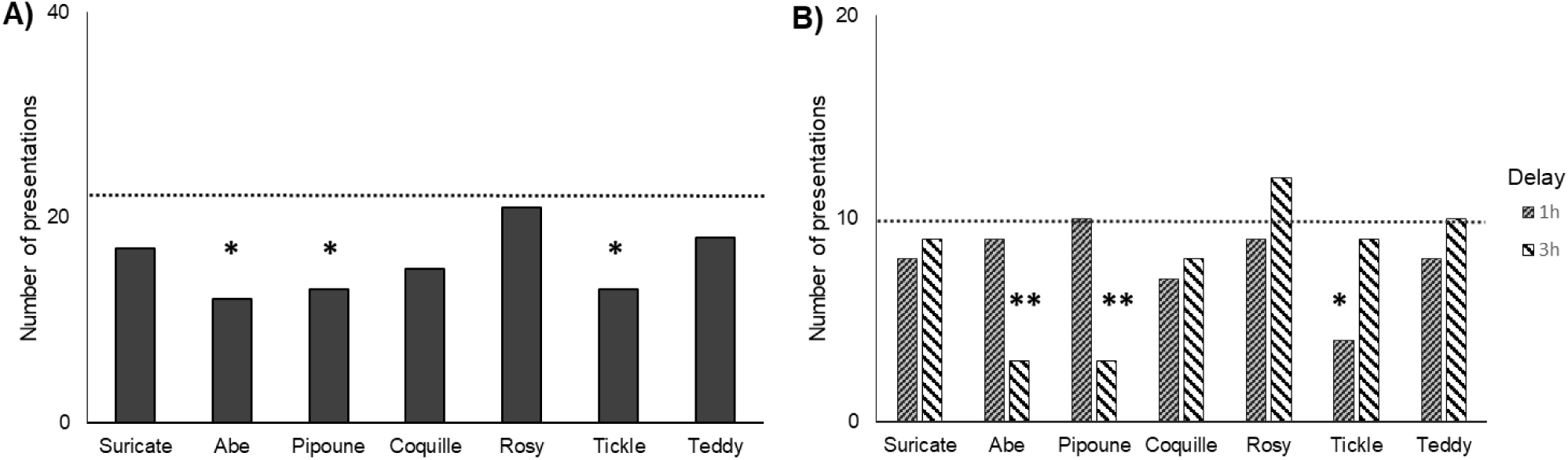
Use of familiarity by each individual during the replenishing rate training. A) Use of familiarity throughout the training. B) Use of familiarity depending on the delay (one hour or three hours) before the second presentation. Asterisks represent a significant avoidance of familiarity (dotted line: chance level; binomial test, * p<0.05, ** p<0.01).

#### Spontaneous alternation

Spontaneous alternation was observed when octopuses alternated their choice between pots of the second presentation between each trial (Fig. 4B)

Considering all trials of all individuals, octopuses did not significantly alternate between pots (114 alternations out of 272 presentations, binomial test, p=0.009), showing constancy rather than alternation in their choices.

When distinguishing the first and the last 20 trials of training, octopuses were significantly averse to alternation during the last 20 trials (54/133, binomial test, p=0.037) but not during the first 20 trials (61/140, binomial test, p=0.151). However, the comparison between the first and the last 20 trials was not significant (two-tailed exact Fisher test, p= 0.842).

At the individual level, five out of seven octopuses almost consistently chose the same pot at one point of training (nine times out of ten consecutive trials; binomial test, p<0.021). The two exceptions were Tickle, showing neither consistency nor alternation, and Coquille, which, contrary to the others, showed a sequence of ten successive alternations (binomial test, p=0.002) during the second half of training.

#### Risk sensitivity

Risk sensitivity appeared in our experiment when subjects chose preferentially one prey over the other during the second presentation of a trial. During this presentation, the less-preferred prey was always available no matter the delay, whereas the preferred food was available half of the time: absent after a delay of one hour, and present after a delay of three hours. Thus, in the absence of delay detection or in the absence of understanding of the delay’s influence, choosing the less-preferred prey was less risky, since it was always available, whereas choosing the preferred prey was riskier, since it was available half of the time (Fig. 4C)

Considering all trials of all individuals, octopuses did not seem to show a preference for the less or more risky option (154 choices of the risky option out of 280 presentations; binomial test, p=0.107).

When distinguishing the first and the last 20 trials of training, octopuses were significantly risk-prone during the first 20 trials (87/140, binomial test, p=0.005), but not during the last 20 trials (67/140, binomial test, p=0.673). However, the comparison between the first and the last 20 trials was not significant (two-tailed exact Fisher test, p=0.228).

At the individual level, a high variability in the response was observed. Three individuals were risk-sensitive throughout training: two individuals (Abe and Teddy) were risk-prone (respectively 31/40 - 27/40; binomial test, p<0.038), whereas one individual (Suricate) was risk-averse (10/40; binomial test, p=0.002; Fig. 6A). Five individuals almost consistently showed either risk-proneness (Abe, Pipoune, Teddy) or risk-aversion (Suricate, Tickle) by choosing the same prey nine times out of ten consecutive trials at one point of training (binomial test, p<0.021).

**Figure 6:**
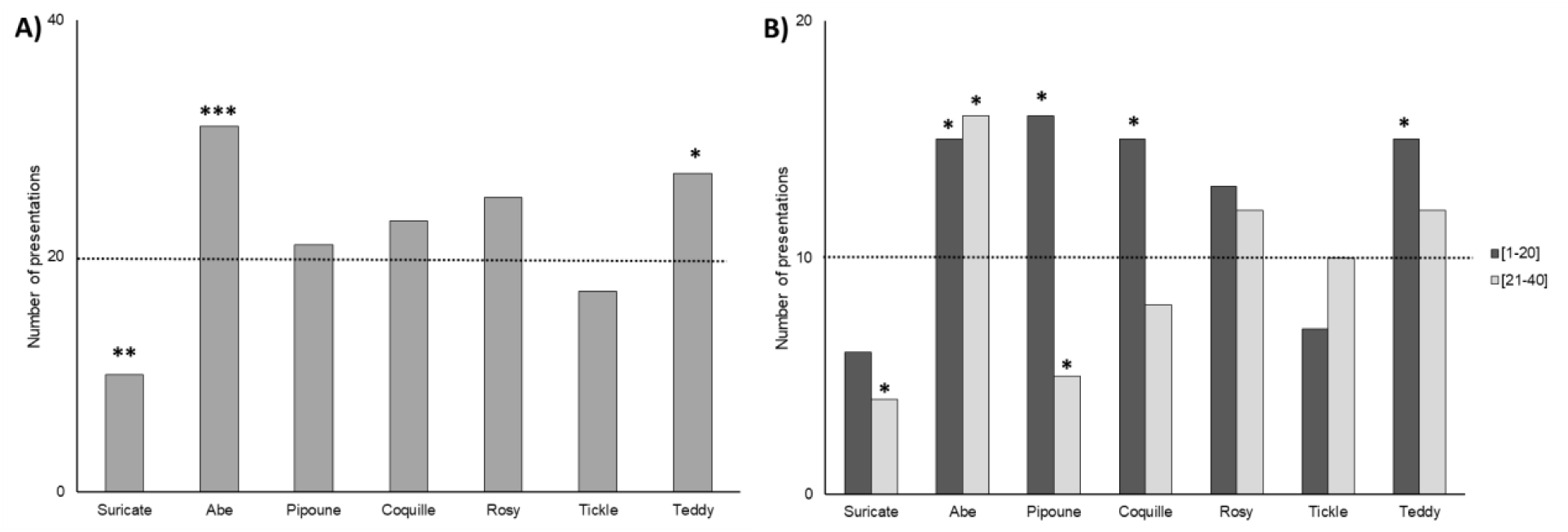
Risk-proneness of each individual throughout the replenishing rate training. A) Risk-proneness throughout the training B) Risk-proneness throughout the training divided between the first and the last 20 trials. Asterisks represent the significant risk-proneness of each individual (dotted line: chance level; binomial test, * p<0.05, ** p<0.01, *** p<0.001).

When distinguishing the first and the last 20 trials of training for each individual, four octopuses (Coquille, Abe, Pipoune, Teddy) were risk-prone during the first training trials (respectively 15/20 - 15/20 - 16/20 - 15/20; binomial test, p<0.042). Interestingly, two of them (Coquille and Teddy) did not show risk-sensitivity anymore during the last 20 trials (respectively 8/20 - 12/20, binomial test, p=0.503), whereas one individual (Abe) continued to be risk-prone (16/20, binomial test, p=0.012), and the last individual (Pipoune) became risk-averse (5/20; binomial test, p=0.041). One other individual (Suricate) showed risk-aversion only in the second half of the training (4/20; binomial test, p=0.012; Fig. 6B).

It is important to note that the preference for one prey over the other was not significant for the first pot opened during the first presentation (before the delay), either at the group level (149/280; binomial test, p=0.310) or at the individual level, except for one individual, namely Abe (29/40, binomial test, p=0.006).

#### Win-stay/Win-shift strategies

Win-shift-Lose-stay strategy could be observed in the second presentation after any delay, when subjects which won (obtained food in the chosen pot) shifted to the other pot the next trial, but when they lost (chose the empty pot) they returned (stayed) on the same pot the next trial. The opposite strategy, Win-stay-Lose-shift, could also be observed when individual which won returned to the same pot on the next trial but changed the chosen pot on the next trial after losing (choosing the empty pot). Since instances of “Lose” were statistically scarce (1/4^th^ of the trials if subjects chose by chance), we focused on “Win” trials during our analyses (Fig. 4D).

Considering all trials of all individuals, octopuses significantly used Win-stay rather than Win-shift strategy (118 choices consistent with Win-stay strategy out of 198 “Win” presentations; binomial test, p=0.008).

When distinguishing the first and the last 20 trials of training, octopuses significantly used Win-stay rather than Win-shift strategy during the “Win” trials in the last 20 trials (64/102, binomial test, p=0.013) but not during the first 20 trials (52/97, binomial test, p=0.543). However, the comparison between the “Win” trials in the first and the last 20 trials of training was not significant (two-tailed exact Fisher test, p=0.559).

At the individual level, two individuals (Suricate and Teddy) significantly used Win-stay rather than Win-shift strategy (respectively 24/35 - 19/26; binomial test, p<0.041; Fig. 7A). When distinguishing the first and the last 20 trials of training for each individual, only Suricate significantly used Win-stay rather than Win-shift strategy in the last 20 trials (14/18; binomial test, p=0.031; Fig. 7B).

**Figure 7:**
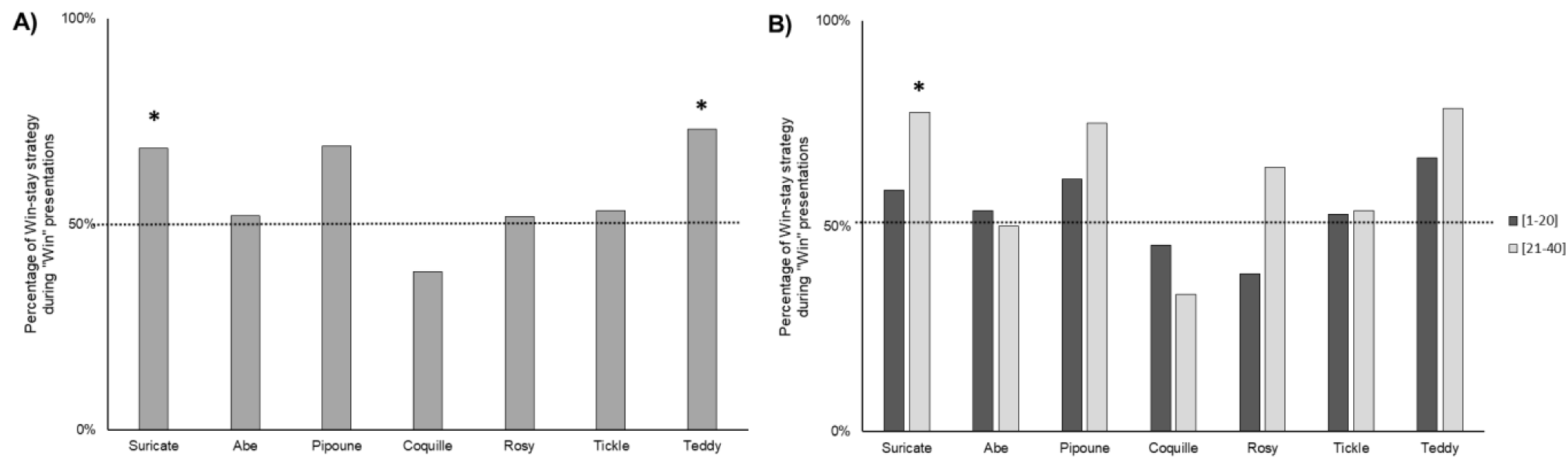
Use of Win-stay strategy by each individual throughout the “Win” presentations of the replenishing rate training. A) Use of Win-stay strategy throughout every “Win” presentations of training. B) Use of Win-stay strategy throughout every “Win” presentations of the training divided between the first and the last 20 trials. Asterisks represent the preferential use of Win-stay strategy (dotted line: chance level; binomial test, * p<0.05).

## Discussion

In our study, seven common octopus (*Octopus vulgaris*) were tested in a task requiring them to keep track of time with different food sources varying in space and time. Most octopuses (six out of seven) relied on less-cognitively demanding strategies than keeping track of time during the replenishing rate learning task. Only one octopus learnt the replenishing rates of different prey types and was able to use these rules to solve an episodic-like memory task.

We analysed the strategies used by tested octopuses during the replenishing rate training: in the beginning, they showed a general risk-proneness, then they tended to avoid the familiar pot and favour constancy in their choices, choosing the same pot over trials, especially when the pot was rewarded. At the individual level, a great variability was observed, with each octopus using and combining strategies in a different way from others.

### Replenishing rate learning and episodic-like memory task

We conducted this experiment to test whether octopuses would favour simpler foraging strategy instead of keeping track of time to solve the task, contrary to cuttlefish which favoured the use of episodic-like memory abilities in a similar experiment (Jozet-Alves et al., 2013). As we hypothesized, most octopuses relied on foraging strategies such as avoidance of familiarity, risk-sensitivity or constancy. Only one individual (Teddy) learnt the replenishing rate of the different food items and subsequently succeeded in the episodic-like memory task.

We can emit several hypotheses to explain why octopuses did not use time tracking in this task while cuttlefish did. Octopuses can be considered in the wild as generalist foragers (Anderson et al., 2008; Mather et al., 2012), with their diet significantly constituted of sessile preys. In the wild, the natural replenishing rates of sessile preys are most probably longer than the time octopuses stay in the same area. It might be needless to remember specifically the time each location was visited, since simple avoidance of food-depleted locations is sufficient.

Another possibility is that octopuses do not spontaneously encode time in temporal distances (i.e. “how long ago it happened”) or with a low accuracy: they may not, or hardly, perceive the time flow as we do, and they may not detect the difference of elapsed delays of one or three hours. This is also the case for rhesus monkeys (Hampton et al., 2005), which cannot distinguish between one hour and 25 hours delays. However, octopuses might encode of time in terms of temporal location (i.e. “when it happened”[within a temporal pattern]) or temporal order (i.e. “before/after what it happened”; (Friedman, 1993; Friedman, 2007). In this case, it would be interesting to study time perception in octopuses, to observe if octopuses really do not keep track of elapsed time, or if our experiment could not bring to light this ability.

If octopuses are able to discriminate different delays, we can wonder why only one individual successfully learnt the replenishing rate task. First of all, we may not have conducted enough trials to make octopuses able to learn the replenishing rates. However, this hypothesis is unlikely: we conducted two to four times more trials (i.e. 40 to 80) than needed by cuttlefish to learn the replenishing rates in previous studies (i.e. 20 trials on average in Jozet-Alves et al., 2013 and Schnell et al., 2021), and the octopus called Teddy needed 43 trials precisely. The three other octopuses trained for 80 trials were given twice the number of trials needed by Teddy to learn without showing any signs of replenishing rates learning.

Another fact that may have hindered the motivation to learn the replenishing rates is that mussels were available at all times in the tank. It might have lowered the pressure of finding food, thus favouring random and simpler foraging strategies. However, we expected octopuses to have a strong hedonic motivation for their preferred food which would have stimulated them to learn its replenishing rate. This belief came from the behaviour observed in cuttlefish: when they know that their preferred food will be ensured at the end of the day, they refrain from eating a less-preferred food available at all times (Billard et al., 2020a). We could extrapolate from this behaviour that cuttlefish are more prey selective than octopuses, however cuttlefish can be generalist in conditions of uncertainty, as they will consume any food during the day when the availability of their preferred food cannot be ensured (Billard et al., 2020a).

Nevertheless, we should not forget Teddy which succeeded on the task. It may indicate that *Octopus vulgaris* might possess the neural prerequisites for episodic-like memory. Cuttlefish and octopuses are phylogenetically close species, both part of neocoleoidea. Both possess a central nervous system, and they share similar brain shape and structures, notably the vertical lobe which is thought to be the place of higher cognitive functions such as memory (Shigeno et al., 2018). However, it should be noted that minor neuroanatomical differences observed between different species of the octopodiform superorder have been mostly linked with the ecology and behaviour of the species (Chung et al., 2022). Sociality notably impacted the formation of gyri in the vertical lobe, indicating that the social differences in octopuses and cuttlefish (*i*.*e*. mostly solitary *vs* socially tolerant) may impact significantly their cognitive abilities.

At last, we should not rule out the use by Teddy of simpler cognitive strategies to solve the task: use of a semantic rule rather than episodic-like memory abilities. Indeed, after a delay of one hour, the memory trace of the first presentation is stronger than after a delay of three hours. Only encoding the position of the pot containing the preferred food and using a simple rule would be efficient to solve the task: when the memory trace of the first presentation is strong, the familiar pot should be avoided, whereas when the memory trace is weak, the familiar pot should be favoured. In this case, only familiarity and semantic learning would have been needed, ruling out episodic-like memory. This hypothesis seems consistent with the fact some individuals chose significantly more the less-familiar pot after a peculiar delay (see below).

### Foraging strategies

Octopuses seem to extensively use familiarity recognition to drive their choices. Instead of choosing the most recently visited pot, most octopuses preferred to go to the least familiar one. In other words, the same pot was first chosen during both presentations of trial s. This can also be considered as an alternation between pots visited during the same trial: octopuses went to the pot A, then B, during the first presentation, then they went back to A during the second presentation. This strategy appeared to be delay-dependent in some individuals: one octopus chose the least familiar pot after a short but not after a long delay, whereas two other individuals presented the reversed pattern. When this strategy is used after an hour but not after three hours, we could hypothesize that the feeling of familiarity faded away progressively, being still present after a one-hourdelay, but not after a three-hour delay. However, the behaviour of the two other individuals seems to contradict this hypothesis, since they chose the pot by avoiding the most familiar one only after a long delay, indicating that the feeling of familiarity might last at least three hours. They could have used familiarity when the memory trace of the first presentation was the weakest (*i*.*e*. after a long delay) and favoured other strategies when the memory trace was stronger (*i*.*e*. after a short delay).

Most octopuses did not present spontaneous alternation behaviour: they significantly kept choosing the same pot trial after trial. There was nevertheless one exception, an octopus which showed significant alternation in its choices at the second presentation from one trial to another. These results are completely to the opposite of what is commonly observed in the literature. Indeed, studies in the wild indicated that octopuses hunt in different areas of their home range on subsequent days (Mather, 1991), since spontaneous alternation is often advantageous during exploration and foraging (Ramey et al., 2009). Moreover, Richman and its colleagues (1986) observed that spontaneous alternation and win-stay strategies seem to be more frequent in low drive vertebrates exposed to small rewards. In the present study, mussels were provided *ad libitum* and small pieces of food were used as rewards, consequently octopuses should have been under low drive, exposed to small rewards, but they did not favour an alternation strategy. Maybe the artificial conditions in which the experiment was conducted incited octopuses to use constancy, but this behaviour stays puzzling.

Spontaneous alternation is often studied in pair with win-shift or win-stay strategies. Indeed, win-shift can be seen as spontaneous alternation if the animal foraging find food each time it visits a new location. The fact that we observed a tendency to use win-stay strategy at the group level is correlated with our observation of alternation avoidance. Indeed, three pots out of four were rewarded on average, so octopuses were rewarded (“win”) most of the time, and since they spontaneously kept going to the same pot, win-stay strategy was observed. However, at the individual level, only two octopuses appeared to rely on this strategy. The behaviour of Suricate can be simply explained by his preference to go for the less-preferred prey (not alternating), which was always available (always “win”). For the other individual, Teddy, such an explanation cannot be used, since it tended to go to the preferred prey, and thus was only rewarded after the three-hours delays. As stated before,

Richman and its colleagues (1986) observed that win-stay strategies are preferentially used by animals with high drive states faced with large rewards. Teddy was the biggest individual out of the seven octopuses, thus we can expect higher needs than the others, which could explain why it relied on a win-stay strategy.

Concerning risk-proneness, high variability was observed. At the group level, but also at the individual level for half of the individuals, octopuses were more risk-prone during the first 20 trials of the training. Two individual showed risk-aversion, and one particular individual, the octopus Pipoune, changed from risk-proneness (first 20 trials of training) to risk-aversion for the subsequent training trials. Empirical studies on risk-sensitivity in vertebrates indicate that when risk come from the variability in the amount or presence of reward, animals are most frequently risk-averse or risk-indifferent, with an influence of the energy budget (Kacelnik and Bateson, 1996). At first sight, our results with octopuses seem to show the contrary. However, we can consider that octopuses required ten to twenty trials to learn that the preferred food is riskier than the less-preferred food. In that case, octopuses presented at first a preference for the pot containing the preferred food. After learning its riskiness, they shifted to risk-indifference or risk-aversion (e.g. Pipoune), or favoured other strategies such as constancy, win-stay, avoidance of familiarity, and for one individual, replenishing rate learning. One individual, Abe, conserved its risk-proneness throughout the training, not fitting into the previous explanation. It is important to add that this individual was the only one to show a preference for one pot (the pot associated to the preferred food) during the first presentation, suggesting a stronger food preference compared to others. It may be explained by the fact that the less-preferred food used for this individual was shelled mussel, even though mussels were always available in the tank. Another possibility is that this highest preference was linked to the position of the pot inside the tank. In our experiment, risk-proneness and position preference could not be distinguished from one another (*i*.*e*. the risky pot was always in the same location). Position preference is a common bias in animals which spontaneously prefer a location of their enclosure. Octopuses possess an arm and an eye preference (Byrne et al., 2002; Byrne et al., 2006a; Byrne et al., 2006b; Frasnelli et al., 2019) which could influence their preference for a peculiar location.

However, since octopuses moved unfearingly in their tank during experiment and their initial position at the beginning of the trial often changed, we focused on risk-proneness rather than position preference.

Another point of attention must be drawn to how individuals combined strategies during training. As stated before, our task created a correlation between constancy and win-stay strategy. Whereas most octopuses used two to three strategies during training, one individual stands out from the others, Rosy, which relied mostly on random choices. Inter-individual differences do not seem to be explained by any physiological or behavioural parameters: sex, size, batch or food preference. Interestingly, in the wild, octopuses individually vary in their prey selection: some are generalists, eating any prey they can find, whereas others are specialists, choosing only one or few types of preys to eat (Mather et al., 2012). This variability cannot be explained by environmental variables only, and individual personality seem to significantly impact octopuses’ behaviour, similarly to our study.

### Conclusion

To investigate episodic-like memory in cephalopods, we reproduced an episodic-like memory task conducted with common cuttlefish, using instead common octopuses. Subjects were asked to keep track of time with different food sources varying in space and time. Contrary to the studies previously undertaken in cuttlefish, most octopuses did not rely on replenishing rates to solve the task, and they favoured simple foraging strategies such as familiarity, constancy, win-stay strategy and risk-sensitivity. One octopus succeeded on the task, likely using episodic-like memory, indicating that octopuses may possess neural substrates for episodic-like memory but may not need to rely on it as strongly as cuttlefish. We hypothesize that the preferential use of different foraging strategies is linked with ecological differences between cuttlefish and octopuses. These results offer new insights on how the ecological intelligence hypothesis and the predatory pressure may shape the use of complex cognitive abilities such as episodic-like memory.

## Acknowledgements

We would like to thank Nicola S. Clayton and Alexandra K. Schnell for their comments on the experimental design, as well as the staff of the marine station (CREC, Unicaen, France) for their help.

## Author contributions

C.J.A., C.B. and L.P designed the studies. L.P and C.D collected the data and ran the analyses.

L.P. wrote the article. All authors discussed the results, revised the manuscript and gave final approval of the version to be published and agree to be held accountable for its content.

## Data availability

Data are available on figshare.com: DOI: 10.6084/m9.figshare.19249079

